# Differential Effects of the Menstrual Cycle on Reactive and Proactive Aggression in Borderline Personality Disorder

**DOI:** 10.1101/452607

**Authors:** Jessica R. Peters, Sarah A. Owens, Katja M. Schmalenberger, Tory A. Eisenlohr-Moul

## Abstract

Borderline personality disorder (BPD) is characterized by rapidly shifting symptoms, including intense anger and aggressive behavior. Understanding how fluctuations in ovarian hormones across the menstrual cycle may contribute to symptom instability is key for accurate assessment of BPD symptoms and effective interventions. Reactive and proactive aggression, as well as anger in and out, were assessed daily in 15 physically healthy, unmedicated naturally cycling female individuals without dysmenorrhea meeting criteria for BPD across 35 days. Urine LH surge and salivary progesterone were used to confirm ovulation and verify cycle phase. Cyclical worsening of symptoms was evaluated using multilevel models to evaluate symptom differences between cycle phases. Both forms of aggressive behavior demonstrated marked cycle effects, with reactive aggression highest during perimenstrual cycle phases, co-occurring with increases in anger in and out. In contrast, highest levels of proactive aggression were observed during the follicular and ovulatory phases, when emotional symptoms and anger were otherwise at lowest levels. These findings highlight the importance of identifying the function of aggression when considering potential psychological and biological influences. Naturally cycling individuals with BPD may be at elevated risk for perimenstrual worsening of a range of interpersonally reactive symptoms, including reactive aggression, whereas proactive aggression may occur more in phases characterized by less emotional and cognitive vulnerability and greater reward sensitivity. Research on aggression in this population should consider cycle effects. Cycling individuals with BPD attempting to reduce aggressive behavior may benefit from cycle-tracking to increase awareness of these effects and to develop appropriate strategies.

## Differential Effects of the Menstrual Cycle on Reactive and Proactive Aggression in Borderline Personality Disorder

Borderline personality disorder (BPD) is a chronic and severe mental condition characterized by rapidly shifting emotional, interpersonal, and behavioral symptoms, including intense anger and aggressive behavior (American Psychiatric Association, 2013). The lability of these symptoms can make aggressive behavior in BPD hard to predict and subsequently more challenging for individuals with BPD, as well as their clinicians and loved ones, to manage. Understanding factors contributing to within-person variability in anger and aggression in BPD is key to improving predictive models of the disorder and optimizing interventions. Given that ovarian steroid hormones estradiol (E2) and progesterone (P4) directly engage neural systems broadly implicated in emotional disorders (Schiller, Johnson, Abate, Schmidt, & Rubinow, 2016), the hormonal changes that occur across the menstrual cycle could provide one source of fluctuating biological vulnerability. We explore the menstrual cycle as a potential source of systematic within-person variability in aggressive behavior for female or assigned female at birth individuals (referred to in this manuscript as cycling individuals) with BPD.

### Aggression in Borderline Personality Disorder

Increased rates of aggression in BPD have been consistently demonstrated across many studies using a wide range of self-report measures, semi-structured interviews, and behavioral tasks (Mancke, Herpertz, & Bertsch, 2015). Aggression in BPD is typically characterized as a response to intense affect, with numerous studies connecting aggressive behavior with the emotion dysregulation, affective reactivity, rejection sensitivity, and overwhelming anger endemic to the disorder (Berenson, Downey, Rafaeli, Coifman, & Paquin, 2011; Mancke, Herpertz, Kleindienst, & Bertsch, 2017; Newhill, Eack, & Mulvey, 2012; Scott, Stepp, & Pilkonis, 2014; Scott et al., 2017). This suggests aggressive behavior in BPD may primarily take the form of reactive aggression, defined as impulsive or uncontrolled aggressive outbursts in response to provocation or frustration (Poulin & Boivin, 2000). In contrast, proactive aggression, defined as premeditated or planned aggression to further one’s goals (Poulin & Boivin, 2000), is more typically conceptualized and observed as a component of antisocial personality disorder (Ross & Babcock, 2009). However, both reactive aggression and proactive aggression have been linked to BPD features in adolescents (Ostrov & Houston, 2008) and adults (Gardner, Archer, & Jackson, 2012). Different mechanisms may underlie these associations, with reactive aggression emerging out of maladaptive coping mechanisms that intensify negative affect, and proactive aggression reflecting more avoidant coping mechanisms (Gardner et al., 2012). Reactive and proactive aggression have also been associated with different facets of impulsivity, with reactive aggression linked to impulsive urges during acute negative affective states and proactive aggression associated with impulsive urges under conditions of heightened positive affect (Hecht & Latzman, 2015). However, much of the research on all types of aggressive behavior in clinical samples with BPD focuses on between-person differences, with more work needed examining factors influencing within-person variability in every day aggressive behavior in BPD (Scott & Pilkonis, 2015).

Limited work to date has studied these within-person fluctuations. One study utilized ecological momentary assessment to examine within-person affective processes contributing to aggression within a community sample of women with recent histories of aggressive behavior (Scott et al., 2017). Those with higher BPD features exhibited a pattern of anger-reactivity to interpersonal rejection leading to aggressive behavior; however, it was not examined whether any physiological factors might influence or exacerbate this pattern of reactive aggression. Identifying these moderators would provide further insight into potential biological and behavioral targets for intervention.

While findings in general population samples demonstrate higher levels of aggression in male participants than female (Knight, Guthrie, Page, & Fabes, 2002), this sex effect is attenuated in most BPD studies, with female participants equally aggressive as male (Mancke, 2015). Despite similar rates of aggressive behavior, some functional neuroimaging findings examining correlates of aggressive behavior demonstrate effects such as diminished striatal serotonin responsivity (Perez-Rodriguez et al., 2012) and deficits in top-down regulatory processes (Herpertz et al., 2017) for male and not female participants with BPD. One possible explanation for fewer distinct physiological correlates found in female participants is that these studies did not control for or examine menstrual cycle effects, which could dramatically increase variability in aggressive behavior and related physiological processes in cycling individuals with BPD, thereby making effects much harder to detect.

### Menstrual Cycle Effects as a Potential Influence

The broader literature on the DSM-5 diagnosis of premenstrual dysphoric disorder (PMDD) demonstrates that hormonal sensitivity is extremely variable across people, with only a subset of cycling individuals (those with PMDD) demonstrating an abnormal emotional and behavioral reactivity to normal cyclical hormone changes (Gehlert, Song, Chang, & Hartlage, 2009; Hartlage, Brandenburg, & Kravitz, 2004; Schmidt, Nieman, Danaceau, Adams, & Rubinow, 1998). It is therefore unsurprising that within unselected samples of cycling individuals, findings of cycle effects on aggression are inconsistent and often null, given that the majority of studies fail to model both the essential *within-person* effects of the cycle or hormone changes and *individual differences* in reactivity to these shifts (see Denson, O’Dean, Blake, & Beames, 2018 for review of such studies). However, within-person research conducted in at-risk samples with PMDD strongly suggests a possibility for cycle effects on aggressive behavior within these particular individuals. Prospective studies have highlighted premenstrual increases in anger and irritability in individuals with PMDD (Landén, Nissbrandt, Allgulander, Sörvik, Ysander, & Eriksson, 2006a), as well as premenstrual deficits in inhibitory control and emotion regulation (Amin, Epperson, Constable, & Canli, 2006; Colzato, Pratt, & Hommel, 2012; Diekhof, 2015; Protopopescu et al., 2005; Wu, Zhou, & Huang, 2014), pointing to the potential for female reproductive hormone fluctuations to influence anger and anger-driven behavior in hormone-sensitive individuals.

Prototypically, a menstrual cycle lasts 28 days, which can be divided into two halves. The *follicular phase* (starting with the onset of menses, lasting until the end of ovulation) is characterized by continuous low levels of P4 and increasing levels of E2 that peak shortly before ovulation. The *luteal phase* (beginning right after ovulation, lasting until the day before the onset of the following menses) is characterized by high, fluctuating levels of both P4 and E2; in the week after ovulation, E2 and P4 rise and peak, and then they fall in the week before menses. Experimental work demonstrates that it is an abnormal sensitivity to this normal luteal hormone flux that triggers mood symptoms in cycling individuals with PMDD (Schmidt et al., 2017).

### Menstrual Cycle Effects in Borderline Personality Disorder

While an early small study had null findings in a sample of psychotropically medicated cycling individuals with BPD (Ziv, Russ, Moline, & Hurt, 1995), more recent work suggests potential menstrual cycle effects on BPD symptom exacerbation. Two studies demonstrated links between reproductive hormone changes and BPD feature expression in non-clinical samples (DeSoto, Geary, Hoard, Sheldon, & Cooper, 2003; Eisenlohr-Moul, DeWall, Girdler, & Segerstrom, 2015). Only one study to date has examined cycle effects on symptoms in a sample (*N*=15) of unmedicated cycling participants diagnosed with BPD (Eisenlohr-Moul et al., 2018). Despite the majority of the sample describing themselves in retrospective reports as experiencing low levels of premenstrual or menstrual symptom changes, in prospective daily assessments, many emotional and behavioral symptoms demonstrated a clear pattern of perimenstrual exacerbation across the sample, with high arousal symptoms such as irritability and anger rising in the luteal phase and peaking in the perimenstrual phase. Lower arousal symptoms, such as depression and hopelessness, were elevated in the perimenstrual phase and extended into the follicular phase. All symptoms examined were lowest in the ovulatory phase. These shifts were present to a clinically significant extent in all but one participant of the study sample; while preliminary findings from a small sample, these findings are consistent with perimenstrual exacerbation potentially occurring in BPD. However, none of these studies to date have specifically reported on menstrual cycle effects on aggressive behavior in cycling individuals with BPD or BPD features.

### Present Study

The present study uses the same sample of unmedicated cycling individuals with BPD previously described (Eisenlohr-Moul et al., 2018) to examine the prospective impact of cycle phase (in ovulatory, normal menstrual cycles) on daily self-reported anger expression and reactive and proactive aggressive behavior in unmedicated females with BPD (*N* = 15) *not* recruited for perceived perimenstrual symptoms. Although participants collected saliva eight times to verify cycle phase, the present study was not powered to examine hormonal effects, instead focusing on cycle phase. We hypothesized that anger in, anger out, and reactive aggression would demonstrate a pattern of significant perimenstrual exacerbation, in similar patterns to other high arousal, interpersonal symptoms. In contrast, we hypothesized no significant cycle effects on proactive aggression, given that this form of aggression is conceptualized as not driven by affective reactivity.

## Methods

### Procedure

Complete details of recruitment and procedures for this sample are presented in Eisenlohr-Moul et al (2018). Listservs, flyers, and social media advertisements were used to recruit “females with emotional and interpersonal problems that interfere with life”; no materials involved references to the menstrual cycle to avoid self-selection bias. In the case of social media advertisements, which were presented only to those identifying as female in their profile, the ads did not mention that the study was only for females. Interested individuals were prompted to complete an online questionnaire assessing exclusionary criteria including: dysmenorrhea in the past 3 months, menstrual cycle length other than 25-35 days, current pregnancy or breastfeeding, psychotropic medication or OC use, illicit drug use, and chronic medical conditions, and schizophrenia or manic episodes. To screen for BPD criteria, this questionnaire also included the *Personality Diagnostic Questionnaire—Borderline Personality Disorder Subscale* (PDQ; Hyler et al., 1988). Individuals who met inclusion criteria including a PDQ score of 5 out of 9 BPD criteria were contacted to complete a phone screen including an interview version of the PDQ. During the phone interview, those who described substantial “pathology, persistence, and pervasiveness” (First & Gibbon, 1997) for at least 5 out of 9 BPD symptoms were invited for a baseline visit.

During the initial visit, a clinical psychologist interviewed individuals using the SCID-PD (BPD module) to determine if they met BPD criteria. Those who met criteria were given a comprehensive orientation to study protocols and completed baseline questionnaires. Beginning on their next menstrual period start day, participants completed 35 days of daily questionnaires to allow for prospective symptom charting. This was done to prevent retrospective bias in symptom reporting; extant literature has demonstrated little correspondence between retrospective reports and prospective assessment of cycle-related symptoms (e.g., Roy-Byrne et al., 1986). Participants also completed daily urine ovulation testing based on individual cycle length and 8 days of passive drool saliva sampling across the cycle. Each individual was compensated $15 for the initial visit and $100 for the remainder of the study. All procedures were approved by the university institutional review board and conducted between 2015-2016.

### Participants

The online screening questionnaire was completed by 310 individuals; of these, 112 of these met inclusion criteria for phone screening (exclusion due to PDQ score (*n*=143), hormone use (*n*=42), irregular cycles (*n*=6), or medical conditions (*n*=7)). Of those invited for a phone screen, 43 individuals elected to complete the screen, and 31 of these were invited for a baseline visit (exclusion due to subthreshold PDQ examples (*n*=8), hormone use (*n*=2), pregnancy/nursing (*n*=2)). Of those invited to the interview, 18 attended and completed the initial visit, and 17 met SCID-BPD criteria and enrolled. One participant did not ovulate during the data collection, and one participant did not comply with study protocols; therefore, the final sample included 15 medically healthy, unmedicated cycling individuals with BPD. Sample characteristics and demographic information are provided in Table 1. All participants reported a lifetime history of psychotherapy; one was engaged in psychotherapy during data collection.

**Table 1.**
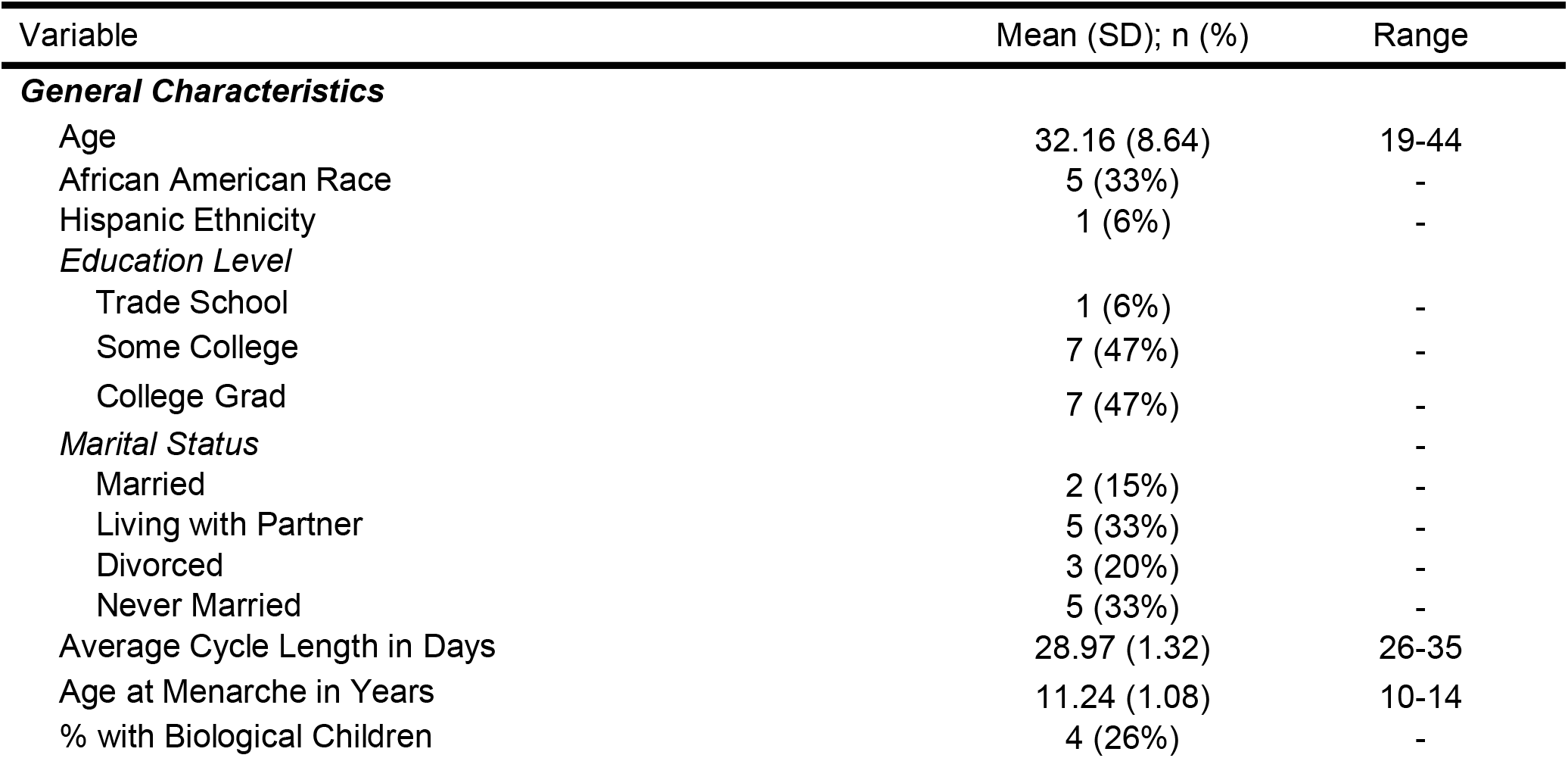
Sample Descriptive Information (*N* = 15)

### Daily Measures

Participants completed daily questionnaires via online survey; links were sent daily via text message and email. Participants completed a range of measures of emotions and behavior; measures of anger and aggression included in the present study are listed below. All items were rated on a response scale from 1=Absent/Not at All to 5=Extreme/Extremely.

***Anger/Irritability*** was assessed using the Daily Record of Severity of Problems (Endicott, Nee, & Harrison, 2006), a daily measure developed to reliably detect menstrually-related changes (e.g. PMDD) in psychiatric and physical symptoms. Only the anger/irritability item is presented in this manuscript.

**Anger In/Out** were assessed with items from the State-Trait Anger Expression Inventory-2 (Spielberger, 1999). *Anger In* was assessed with two items: “I boiled inside but did not show it” and “I was angrier than I was willing to admit.” *Anger Out* was assessed with two items: “I expressed my anger” and “I argued with others”.

***Aggression*** was assessed with items from the Reactive-Proactive Aggression Questionnaire (Raine et al., 2006). *Reactive Aggression* was assessed with the two items: “I yelled at others when they annoyed me” and “I reacted angrily when provoked by others.” *Proactive Aggression* was assessed with the two items: “I yelled at others so they would do things for me” and “I had fights with others to show who was on top.”

### Ovulation Testing and Salivary Ovarian Steroids

Participants were provided urine luteinizing hormone (LH) tests by the brand Clearblue and instructed to test daily starting 7 days after onset of menses to confirm ovulation. With ovulation occurring approximately 12-36 hours after the LH surge, the day after the first positive LH test was set as the ovulation day. This testing kit has been validated against ultrasound and reduces false positives by applying a LH threshold of 40 pg/mL. To assay for E2 and P4 for cycle phase validation, on days 2, 3, 8, and 9 following menstrual onset and following the positive LH test, participants were asked to collect saliva. Polypropylene vials were filled with 5 mL of saliva via passive drool between and stored in a temperature stability pouch in their home freezer; once all samples were collected, the pouch was returned to the laboratory. Based on pilot data suggesting poor compliance in morning saliva collection, participants were asked to collect samples between 4 and 6pm in the afternoon. Hormone assays were completed at the UNC Chapel Hill School of Medicine Core Laboratory with RIA kits by the brand Salimetrics (Carlsbad, CA).

#### Reliability and Validity of Salivary E2 and P4

Intra-assay CVs were 6.3% for E2 and 4.0% for P4, while inter-assay CVs were 8.9% for E2 and 5.5% for P4. Per Salimetrics validation reports, the Salimetrics RIA method sensitivity is .1pg/ml and 5pg/ml for E2 and P4, respectively. Salivary E2 and P4 changes across the cycle should reflect the cyclical changes of serum E2 and P4. In the current study, P4 levels displayed the expected luteal rise in each participant. In contrast, each participant’s salivary E2 levels did not show any of the expected cyclical changes indicating problems with sampling, storage, or assay since ovulation was confirmed by the post-ovulatory rise of P4 levels. It could be hypothesized that the afternoon saliva sampling (with generally reduced E2 levels in the second half of the day compared to the first) in combination with the storage of the saliva in the home freezer might account for these problems. Consequently, only P4 levels were utilized to verify cycle phase.

### Menstrual Cycle Phase Coding

We distinguished four cycle phases—*ovulatory, midluteal, perimenstrual*, and *follicular*— for each participant using several sources of information: (1) forward and backward counting from the day of menstrual onset (Edler, Lipson, & Keel, 2007), (2) ovulation confirmed by a positive LH test result, and (3) the marked rise of P4 levels confirmed by hormone assays. The *ovulatory phase* was set to the days (−15) to (−12) prior to the subsequent onset of menses (backward counting) and validated by a positive LH-based ovulation test and an exponential rise of P4 levels following ovulation. The *perimenstrual phase* was defined as days (−4) to (+3), where there is no day 0 and day +1 marks the onset of menses. To additionally validate this perimenstrual window, the typical length of the luteal half of the cycle (14 plus/minus 2 days) was taken into consideration. The days between the ovulatory and the perimenstrual phases formed the *midluteal phase*, which was validated by high levels of P4. Finally, the days between the perimenstrual and the ovulatory phases were defined as the *follicular phase*, being additionally validated by low levels of P4.

### Data Analysis and Power

To improve interpretability and isolate the within-person component of the outcome, each outcome was person-standardized (as in Edler et al., 2007; Klump et al., 2013). To test the hypotheses, multilevel models evaluated the impact of cycle phase (coded as a categorical variable, with alternating reference phases). SAS PROC MIXED accounted for the nested structure of observations (i.e., random intercept) as well as the autocorrelation of today‘s rating with yesterday’s rating. Random effects of cycle phase contrasts were not included as they did not improve model fit; this indicates that the impact of the menstrual cycle on symptoms were roughly uniform in this sample. Five-day rolling averages of each person-standardized outcome were utilized for graphical depictions.

To determine power for phase contrasts, we calculated the design effect to determine the smallest detectible effect size (Snijders, 2005). Each participant provided up to 35 daily surveys (lower-level *N* was 490), and anger and aggression generally showed low intraclass correlations (i.e., a high degree of flux within a person, average ICC across anger and aggression items = .18). Power for these models was adequate to detect conventionally small-to-medium sized contrasts between phases (average smallest detectible effect size was an *f^2^* of .10). That is, the study was powered to detect significant phase contrasts accounting for at least 10% of the variance in the daily outcome. Age was considered as a covariate but ultimately omitted as it was not associated with outcomes and did not alter our pattern of findings.

## Results

### Descriptives

Table 1 provides demographic and clinical characteristics. The sample was diverse with regard to age and race. Information about psychological symptoms in this sample has been previously published (Eisenlohr-Moul et al., 2018); participants endorsed levels of affective symptoms and of BPD symptoms consistent with their BPD diagnosis. 93% of surveys were completed, and 90% of salivary samples were completed.

### Contrasting Anger and Aggression Across Menstrual Cycle Phases

Results of multilevel models testing phase effects on anger and aggression are presented in Table 2. All variables demonstrated significant cyclical changes; Figures 1-5 depict patterns of change across cycle day and cycle phase for each. Graphs include smoothed, person-standardized P4 values.

**Table 2.** *Multilevel Model Results: Within-Person Menstrual Cycle Phase Contrasts for Aggression-Related Variables (N = 15)*.

|  | **Within-Person Phase Contrasts (Standardized Fixed Effects)** |  |  |  |  |  |
| --- | --- | --- | --- | --- | --- | --- |
|  | Perimenstrual Phase<br>Reference<br>(Falling E/P) |  |  | Midluteal Phase<br>Reference<br>(High Stable E/P) |  | Follicular Phase<br>Reference<br>(Low Stable E/P) |
|  | v. Follicular<br>(Low Stable E/P) | v. Midluteal<br>(High Stable E/P) | v. Ovulatory<br>(Rising E/Low P) | v. Ovulatory<br>(Rising E/Low P) | v. Follicular<br>(Low E/P) | v. Ovulatory<br>(Rising E/Low P) |
| Anger/Irritability | **-.51\*\*\* (.13)** | **-.24\* (.11)** | **-.76\*\*\* (.17)** | **-.52\*\* (.18)** | **-.27\* (.13)** | -.24 (.17) |
| Anger In | .071 (.15) | -.014 (.16) | **-.19\* (.08)** | -.11 (.21) | .086 (.16) | -.12 (.12) |
| Anger Out | -.056 (.16) | .10 (.18) | .14 (.12) | .033 (.22) | -.12 (.13) | .13 (.20) |
| Reactive Aggression | -.10 (.17) | **.27\* (.09)** | **-.22\* (.11)** | **-.26\*\* (.10)** | **-.31\* (.13)** | .04 (.22) |
| Proactive Aggression | **.31\* (.15)** | .10 (.18) | **.45\* (.20)** | **.27\* (.13)** | .15 (.17) | **.23\* (.11)** |

*Note.* \* $p < .05$ , \*\* $p < .01$ , \*\*\* $p < .001$ . Reference means that that phase was set as the reference category in the category coding used in multilevel regression models.

**Figure 1.**
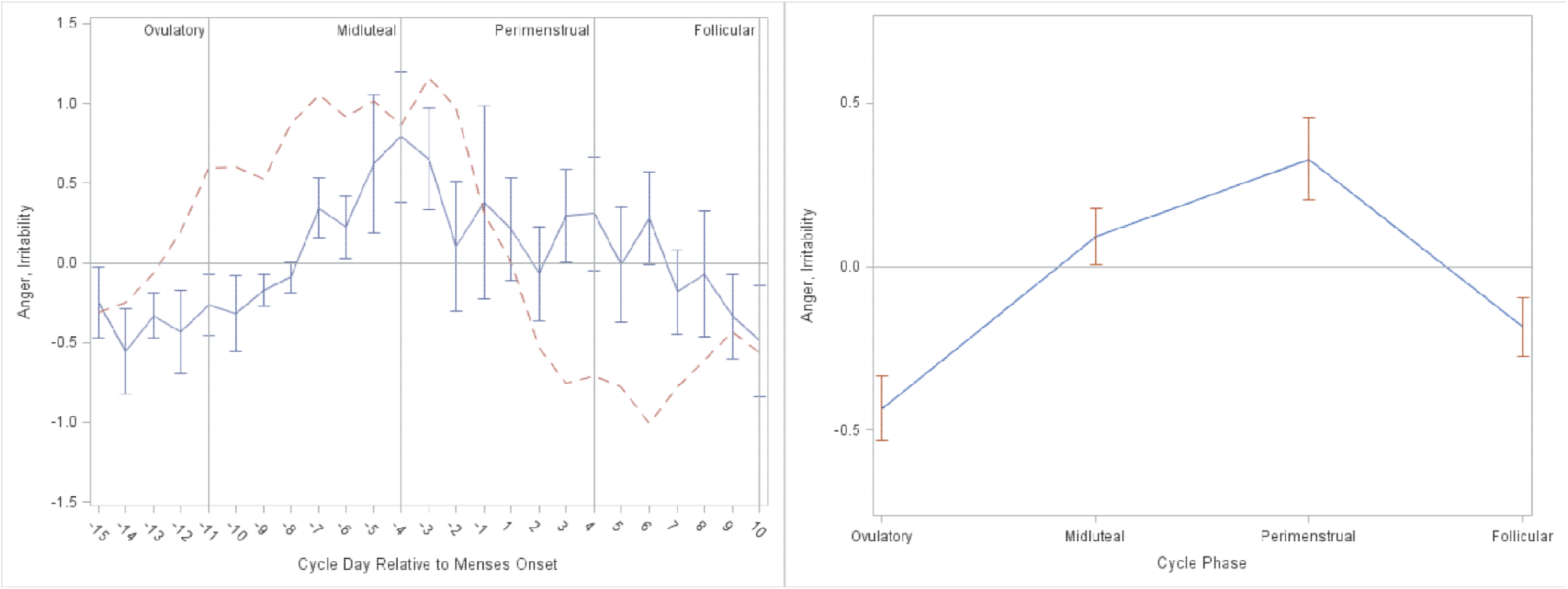
Person-Standardized Progesterone (dashed line) and Anger/Irritability Across Menstrual Cycle Day (Panel A) and Person-Standardized Anger/Irritability Across Cycle Phase (Panel B) in 15 People with BPD.

As reported previously, *Anger/Irritability* began rising during the midluteal phase, during which levels were significantly higher than the follicular and ovulatory phases. *Anger/Irritability* continued to rise into the perimenstrual phase, during which levels were significantly higher than all other phases (see Figure 1). Levels of *Anger-In* increased later in the cycle but were similarly highest in the perimenstrual phase, demonstrating significant contrasts with the midluteal, follicular, and ovulatory phases (see Figure 2). In contrast, *Anger-Out* did not demonstrate any significant cyclical patterns (see Figure 3).

**Figure 2.**
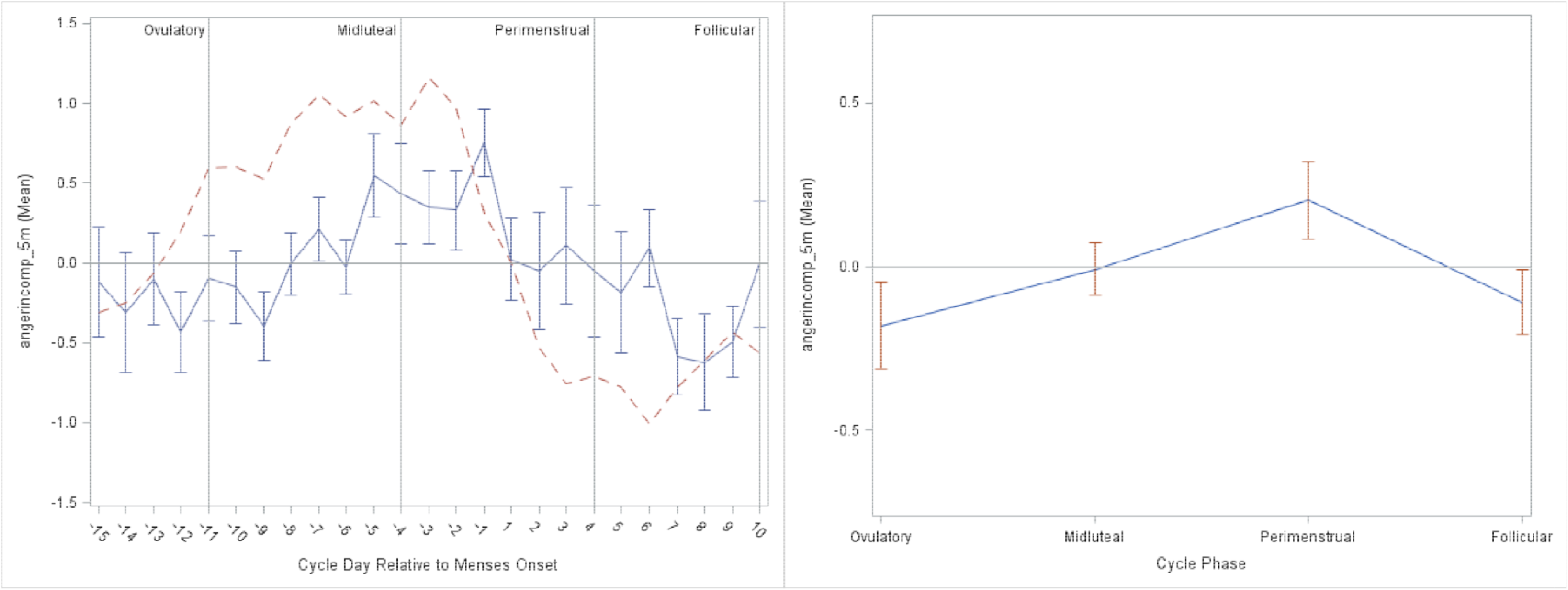
Person-Standardized Progesterone (dashed line) and Anger In Across Menstrual Cycle Day (Panel A) and Person-Standardized Anger In Across Cycle Phase (Panel B) in 15 People with BPD.

**Figure 3.**
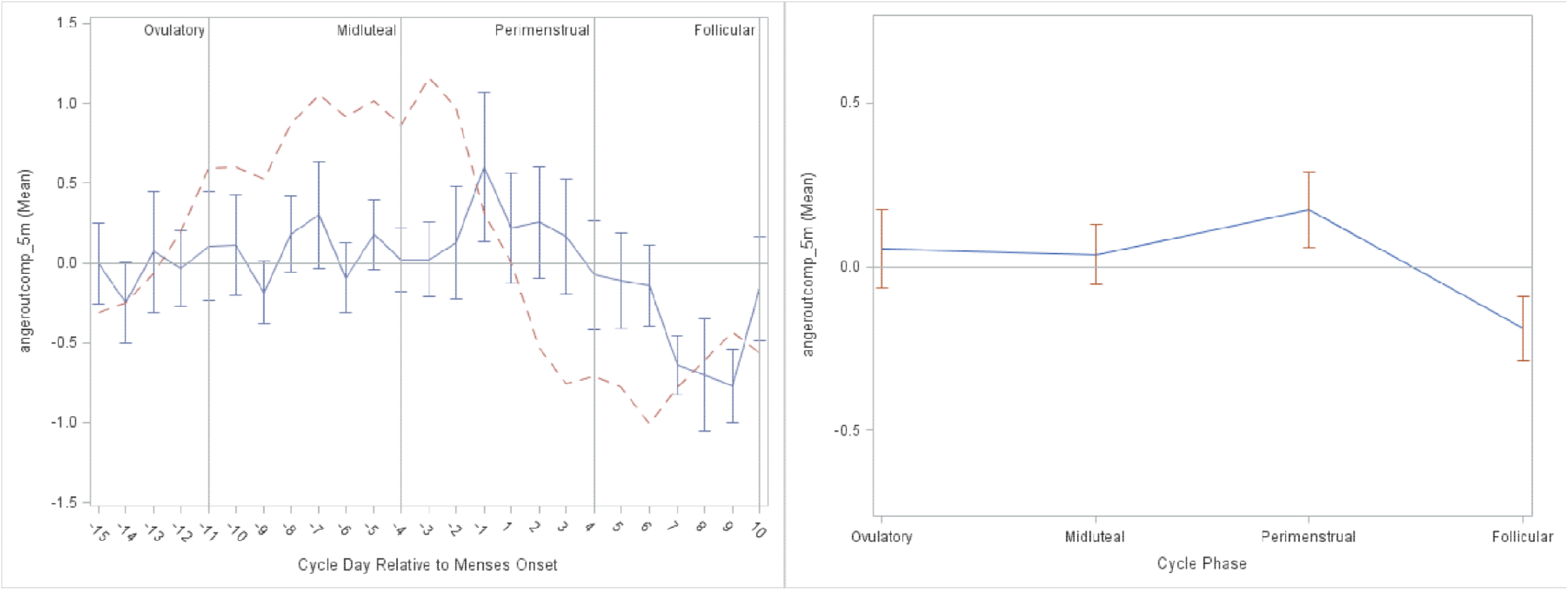
Person-Standardized Progesterone (dashed line) and Anger Out Across Menstrual Cycle Day (Panel A) and Person-Standardized Anger Out Across Cycle Phase (Panel B) in 15 People with BPD.

Levels of reactive aggression were greatest in the midluteal phase, with significant contrasts with all other phases. Reactive aggression in the perimenstrual phase was also elevated relative to the ovulatory phase, when reactive aggression was lowest (see Table 2). This partially supports our prediction, showing that cycle-based exacerbation of reactive aggression occurs, but beginning and peaking in the midluteal phase and then continuing into the perimenstrual phase to a lesser degree (see Figure 4). Of note, these findings for reactive aggression track with the cyclical patterns of anger/irritability, with reactive aggression at highest levels as anger/irritability first rises.

**Figure 4.**
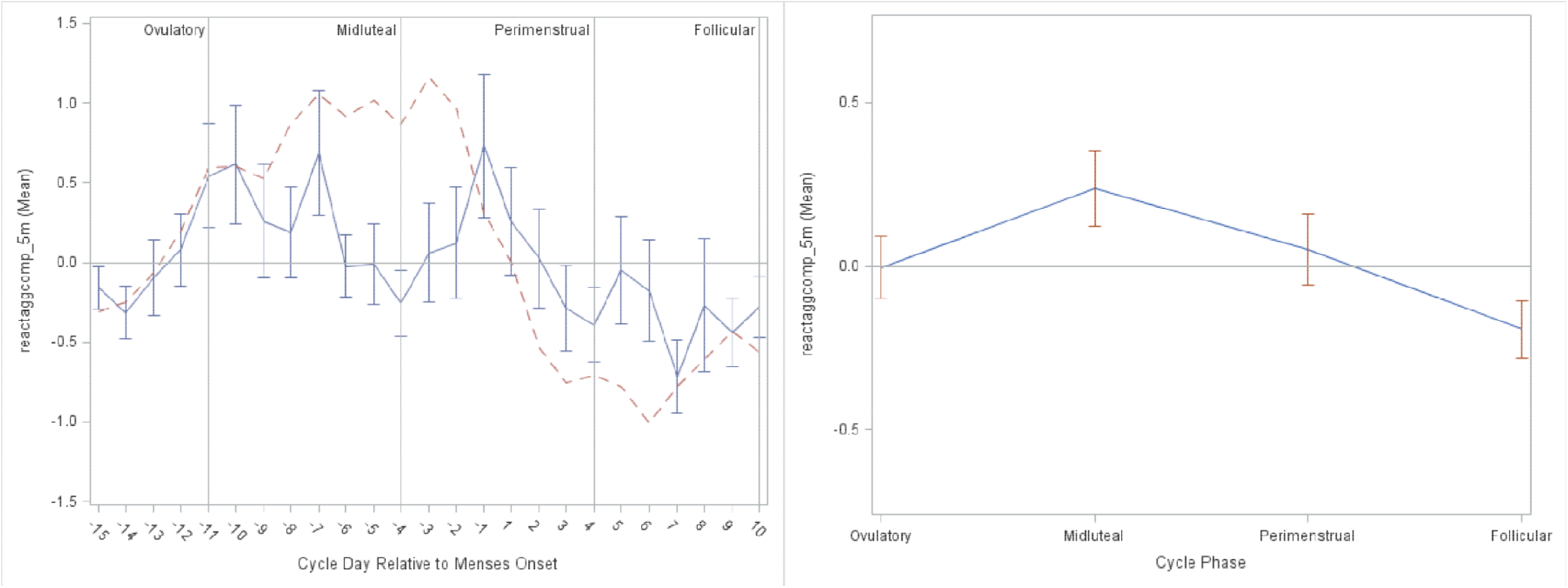
Person-Standardized Progesterone (dashed line) and Reactive Aggression Across Menstrual Cycle Day (Panel A) and Person-Standardized Reactive Aggression Across Cycle Phase (Panel B) in 15 People with BPD.

Our second hypothesis was a lack of cycle effects on proactive aggression. This hypothesis was not supported; instead, proactive aggression was at highest levels during the *ovulatory* phase, relative to the perimenstrual, midluteal, and follicular phases, with lowest levels occurring in the perimenstrual phase (see Figure 5). This unexpected finding represents an opposite pattern to the perimenstrual exacerbation found for most other BPD symptoms and for anger and reactive aggression.

**Figure 5.**
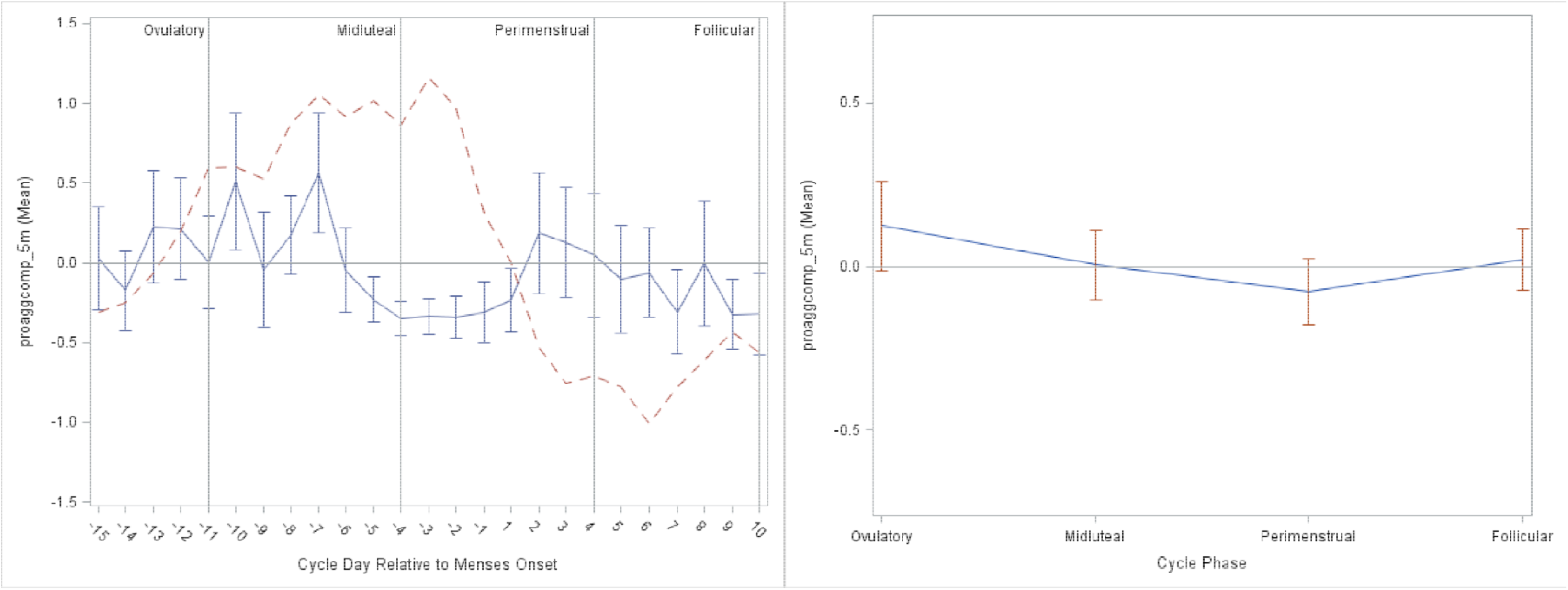
Person-Standardized Progesterone (dashed line) and Proactive Aggression Across Menstrual Cycle Day (Panel A) and Person-Standardized Proactive Aggression Across Cycle Phase (Panel B) in 15 People with BPD.

## Discussion

We examined menstrual cycle effects on anger, anger expression, and aggression in unmedicated cycling individuals with BPD. The observed patterns can be described as follows. First, all forms of anger and aggression generally show lowest levels in the ovulatory phase, with the exception of proactive aggression which was at highest levels. Second, in the midluteal phase, anger and irritability rise, accompanied by peak levels of reactive aggression. Third, in the perimenstrual phase, anger/irritability peak and reactive aggression continues to be relatively high, while anger-in also becomes elevated. Previous findings with this sample have demonstrated this phase to also include heightened anger rumination, felt-invalidation, and rejection sensitivity (Eisenlohr-Moul et al., 2018). Finally, anger and reactive aggression return to baseline in the follicular phase.

These findings highlight the importance of identifying the function of aggression when considering potential psychological and biological influences. Irritability has been identified as a cardinal symptom of PMDD and is thought to arise from abnormal luteal changes in the serotonin system (Landén, Nissbrandt, Allgulander, Sörvik, Ysander, & Eriksson, 2006b; Rapkin et al., 1987), as well as abnormal responses to normal luteal changes in the GABAergic neurosteroid metabolites of P4 (Martinez et al., 2016). Given the similar timing of increased irritability in this sample with BPD, a similar physiological process may underlie increased anger/irritability in the midluteal and perimenstrual phases. This sudden shift in irritability may drive reactive aggression in particular, as it peaks earlier than other interpersonally reactive symptoms (e.g., anger-in, anger rumination, rejection sensitivity, perceived invalidation). It is possible that reactive aggression reflects an immediate impulsive response to the initially heightened irritability, whereas these other symptoms characteristic of BPD build as interpersonal dynamics feel more strained over longer amounts of time.

For those who are sensitive to cyclical hormone fluctuations, increases in irritability may coincide with perimenstrual decreases in effective emotion regulation (Wu et al., 2014) and luteal (vs. mid-to-late follicular) reductions in neural activation in brain regions underlying inhibitory control under conditions of negative affect (Protopopescu et al., 2005; 2008). Reactive aggression (vs. proactive aggression) has been cross-sectionally associated with negative urgency, a propensity towards impulsive action under conditions of heightened negative affect (Hecht & Latzman, 2015). Thus, elevations in reactive aggression during the midluteal and perimenstrual phases among sensitive individuals are likely facilitated by increases in negative affect coupled with reductions in effective emotion regulation and inhibitory control.

In contrast, ovulatory increases in proactive aggression occurred when anger and other symptoms were generally at lowest levels in this sample. While there is considerable inter-individual variability, in most cycling individuals, the ovulatory phase is also associated with less emotional and cognitive vulnerability (Owens & Eisenlohr-Moul, 2018; Schiller et al., 2016). This generally protective effect of higher levels of estradiol may provide individuals with BPD greater ability to engage in premeditated and purposeful (versus reactive) action. These actions, however, may still be influenced by the negative cognitive biases held by individuals with BPD that the world is hostile, dangerous, and untrustworthy and that preemptive steps may be required to protect oneself from being rejected or attacked (see Baer, Peters, Eisenlohr-Moul, Geiger, & Sauer, 2012 for review). Within the context of BPD, greater relative cognitive resources at ovulation may therefore be expressed with higher levels of proactive aggression in attempts to protect the self from anticipated threat or in what is seen as the most viable way to meet needs effectively.

Ovulatory increases in proactive aggression may also be driven by changes in reward sensitivity. While reactive aggression is associated with negative urgency (the behavioral tendency towards impulsive action under conditions of heightened negative affect), proactive aggression is uniquely associated with positive urgency (the behavioral tendency towards impulsive action under conditions of heightened positive affect) in between-person studies (Hecht & Latzman, 2015). Hormone-sensitive individuals demonstrate greater reward-related neural activation during the midfollicular phase as compared to the luteal phase, indicating increased reward sensitivity when E2 is elevated (Bayer, Bandurski, & Sommer, 2013; Dreher et al., 2007). Consistent with this, many appetitive urges like substance craving and sexual desire are also positively associated with E2 levels and are elevated in the follicular phase compared to the luteal phase (Jones et al., 2018; Roney & Simmons, 2013; 2016). Thus, under conditions of elevated E2, heightened neural reactivity to anticipated reward and increased appetitive urges in the follicular and ovulatory phases may elevate susceptibility to engagement in proactive aggression to facilitate reward attainment.

These preliminary findings on cycle effects on aggressive behavior in BPD warrant further research. The present study used a small sample, which limits generalizability; these findings indicate the need to conduct much larger studies of menstrual cycle effects in BPD. Contrasting this group with healthy and clinical controls would clarify whether these patterns are specific to BPD or represent tendencies seen across individuals experiencing cycle effects on psychological functioning. Additionally, using multiple measures of aggressive behavior, including behavioral tasks, would provide more robust evidence.

In conclusion, these findings also suggest the importance of considering menstrual cycle effects when conducting research on and practicing clinical work with cycling individuals with BPD. It may be important to include the menstrual cycle in statistical models. For between-person studies, this may involve controlling for cycle phase (which can be determined for any given day with the dates of the last menstrual period and of the next, which allows for the kind of menstrual cycle phasing done in the present and similar studies, e.g., Edler et al., 2007) or attempting to recruit cycling individuals at the same cycle phase (e.g., five days following the onset of menses, or seven days after a positive ovulation test). Given that reactive aggression is highest in the midluteal and perimenstrual phases, studies may have more power to detect effects or examine mechanisms if all participants are in these phases. For within-person studies, especially daily diary or experience sampling studies, including cycle phase as either a covariate or moderator of effects may reduce noise in the data and clarify effects of interest.

While no study to date has attempted pharmacological intervention to address cyclical BPD symptom exacerbation, selective-serotonin reuptake inhibitors (SSRIs) administered either continuously or intermittently during following ovulation until menstruation have been demonstrated to reduce similar cycle effects specifically on irritability in PMDD (e.g., Cunningham, Yonkers, O’Brien, & Eriksson, 2009; Steiner et al., 2006). RCTs examining efficacy of these medications in reducing cycle-reactivity in individuals with BPD are warranted. In addition to pharmacological approaches, individuals with BPD attempting to reduce aggressive behavior may benefit from cycle-tracking to increase awareness of cycle effects and to develop appropriate strategies. This can be easily integrated into therapies such as Dialectical Behavior Therapy (Linehan, 1993; 2014) that have patients track daily symptoms as an intervention component. Individuals who observe menstrual cycle patterns can then work with their clinical team to develop strategies to more effectively manage changes in irritability and to reduce reactive aggression. These could include, during days of highest reactive aggression risk, proactively utilizing self-soothing to manage feelings of irritability and anger, planning ahead with partners and family to pause and postpone when possible conversations with high potential for conflict, and setting reminders to use skills and therapist coaching. In contrast, ovulatory phases may be ideal times to work on challenging cognitive biases that suggest aggressive behavior is required to meet needs and practicing more effective interpersonal strategies and skills.

## Acknowledgements

This work was supported by grants from the National Institute of Mental Health (K23MH112889, R00MH109667; R01MH099076) and the National Center for Advancing Translational Sciences (UL1TR001111) at the National Institutes of Health.

